# Prefusion-Stabilized Lassa Virus Trimer Identifies Neutralizing Nanobodies and Reveals an Apex-Situated Site of Vulnerability

**DOI:** 10.1101/2022.04.21.488985

**Authors:** Jason Gorman, Crystal Sao-Fong Cheung, Zhijian Duan, Yaping Sun, Pengfei Wang, Jeffrey C. Boyington, Andrea Biju, Tatsiana Bylund, Cheng Cheng, Li Ou, Tyler Stephens, Yaroslav Tsybovsky, Raffaello Verardi, Shuishu Wang, Yongping Yang, Baoshan Zhang, ChengYan Zheng, Tongqing Zhou, John R. Mascola, David D. Ho, Mitchell Ho, Peter D. Kwong

## Abstract

Lassa virus (LASV) is responsible for 100,000-300,000 zoonotic infections annually and poses a threat to public health. Development of antibody-based therapeutics or vaccines has been challenging because neutralizing antibodies – even among Lassa hemorrhagic fever survivors – are generally of low titer, and the target of neutralizing antibodies, the trimeric glycoprotein complex (GPC), a type 1-fusion machine with GP1 and GP2 subunits, has been difficult to produce. Here, we use structure-based design to obtain a soluble LASV GPC by engineering an inter-protomer disulfide (R207GC_GP1_-L326C_GP2_) and appending the T4-fibritin trimerization domain. We verified the antigenicity of this prefusion-stabilized LASV GPC against a panel of human antibodies and used electron microscopy (EM) to confirm its trimeric association. We panned the prefusion-stabilized LASV GPC against single domain ‘nanobody’ libraries and identified one of camel origin, which we named D5, which bound GPC with 27 nM affinity and neutralized the Josiah strain of LASV with an IC_50_ of 12 µg/ml when formatted into a bivalent IgG2a context. The cryo-EM structure of a ternary complex of the D5 nanobody, the antigen-binding fragment of human antibody 8.11G, and LASV GPC revealed D5 to recognize a site-of-vulnerability at the trimer apex. The recognized site appeared to be specific to GPC lacking cleavage of between GP1 and GP2 subunits. Collectively, our findings suggest that GPC-cleavage intermediates may be targets for LASV neutralization and define an apex-situated site of vulnerability for vaccine development.

**Significance:** Lassa virus (LASV) infection is expanding outside its traditionally endemic areas in West Africa, posing a biothreat to the world. LASV-neutralizing antibodies, moreover, have proven difficult to elicit. To gain insight into requirements for antibody-mediated neutralization of LASV, we developed a prefusion-stabilized LASV glycoprotein trimer (GPC), panned it against phage libraries comprised single-domain antibodies or nanobodies from shark and camel, and identified one, D5, which – when placed into bivalent IgG2a context – could neutralize LASV. Cryo-EM analysis revealed D5 to recognize a cleavage-dependent site-of-vulnerability at the trimer apex. We propose this apex-situated site to be an attractive target for LASV vaccine and therapeutic development.

## INTRODUCTION

Lassa virus (LASV) is an enveloped RNA virus, an Old World arenavirus, which causes the acute viral hemorrhagic illness Lassa fever. Lassa fever is prevalent in West Africa, affecting 100,000 to 300,000 of individuals each year (https://www.cdc.gov/vhf/lassa/index.html) and causing approximately 5,000 deaths (1, 2). To date, no licensed vaccine is available for the prevention of Lassa fever, and the only treatment is Ribavirin, a broad spectrum antiviral drug (3). LASV is usually initially spread to people via contact with urine or feces of infected rodents; human-to-human transmission can then occur via direct contact. LASV has been included on the priority pathogen list for WHO’s R&D Blueprint for Action to Prevent Epidemics in an urgent effort to develop effective vaccines (4, 5).

The surface of LASV virions is covered by the trimeric type 1-fusion glycoprotein complex (GPC) (6, 7), the only antigen available for virus-neutralizing antibodies. Each protomer of the GPC consists of a receptor-binding GP1 subunit, a transmembrane-spanning GP2 subunit, and the stably-associated signal peptide (SSP) (8-12). The highly glycosylated GPC induces weak and inconsistent immune responses in both natural infection and vaccination (13-17). Indeed, only 16 LASV-neutralizing antibodies have been reported, which were isolated from 17 Lassa fever convalescent patients by analyzing over 100 antibodies (18). These 16 neutralizing antibodies fall into four competition groups: GP1-A, GPC-A, GPC-B, and GPC-C, based on their recognition and cross-reactivity (18). The GP1-A group comprises three antibodies (10.4B, 12.1F, and 19.7E) that bind GP1 but not GP2. The other three groups recognize only the fully assembled GPC. GPC-A group comprises three antibodies: 8.11G, 25.10C, and 36.1F; GPC-B comprises nine antibodies: 2.9D, 18.5C, 25.6A, 36.9F, 37.2D, 37.2G, 37.7H, and NE13; and GPC-C comprises a single antibody: 8.9F (18). GP1-A and GPC-A antibodies recognize LASV GPC with each Fab binding one single GPC protomer (19). By contrast, each GPC-B Fab binds across two adjacent GPC protomers at the interface in the assembly (8). Such binding that bridges two GPC protomers, referred to as inter-protomer quaternary recognition, effectively locks the trimers in a prefusion state to mediate neutralization (8, 20). The epitope for the GPC-C antibody, 8.9F, remains unclear.

To provide insight into LASV GPC requirements for antibody-mediated neutralization, we designed a stable soluble GPC trimer based on the previously published GPC structure (PDB: 5VK2) (8), by appending a foldon trimerization domain and engineering an inter-protomer disulfide. We verified the prefusion conformation of this stabilized GPC trimer by antigenic and cryo-EM studies. We then used phage display to identify single domain antibodies (also called nanobodies) that bound the stabilized GPC trimer from both the shark variable domain of New Antigen Receptors (V_NAR_) (21, 22) and camel single variable domain from heavy chain-only (V_H_H) antibodies libraries. We placed binding nanobodies into a bivalent IgG2a context to assess neutralization, carried out competition mapping between both neutralizing nanobodies and human antibodies, and determined the cryo-EM structure of the most potent neutralizing nanobody. The revealed nanobody epitope, at the trimer apex, appeared to require a lack of cleavage between GP1 and GP2 subunits. Altogether, this study highlights the development of a soluble prefusion-stabilized LASV trimer, identifies nanobodies capable of neutralizing LASV, and reveals an apex-situated site-of-vulnerability as a vaccine target.

## RESULTS

### Structure-based design and characterization of prefusion-stabilized LASV GPC

As a type I-viral fusion machine, the GPC is metastable and refolds from the metastable prefusion conformation to the more stable post-fusion conformation (6, 23). Since the epitopes for most neutralizing antibodies are present only in the prefusion conformation of the trimer, we sought to stabilize this conformation of the trimer. Prior prefusion stabilizaton by Hastie et al. (8, 20) identified an intra-protomer disulfide as well as a modified cleavage site (GPCysR4); this construct formed monomers, but in the presence of quaternary specific antibodies like 37.7H, 25.6A, or 18.5C, formed GPC trimer. Epitopes of these antibodies overlap, and each involves two adjacent GPC monomers that stablizes the prefusion trimeric conformation. Starting with the structure of the prefusion LASV GPCysR4, we designed over 150 variants (**Supplemental Dataset 1**), which we screened for binding to the quaternary-specific GPC antibody 37.7H. From the screening results (**Fig. S1**), we identified an engineered inter-protomer disulfide bond that links GP1 subunit of one protomer to the GP2 subunit of a neighboring protomer to yield improved antigenticity. This inter-protomer disulfide, R207GC-L326C, replaced the existing intra-protomer disulfide C207-C360 present in GPCysR4 by reverting the residue at position 360 to a Gly and introducing a L326C mutation. An additional Gly residue was inserted after position 206 (G206a) to allow the two Cys side chains to have optimal geometry for the formation of 207C_GP1_-L326C_GP2_ inter-protomer disulfide bond (**Fig. 1A)**. To further stabilize the trimeric conformation of the LASV GPC, we appended a T4-fibritin (foldon) trimerization domain at the C-terminus of GP2 soluble construct (**Fig. 1A)**.

**Fig. 1.**
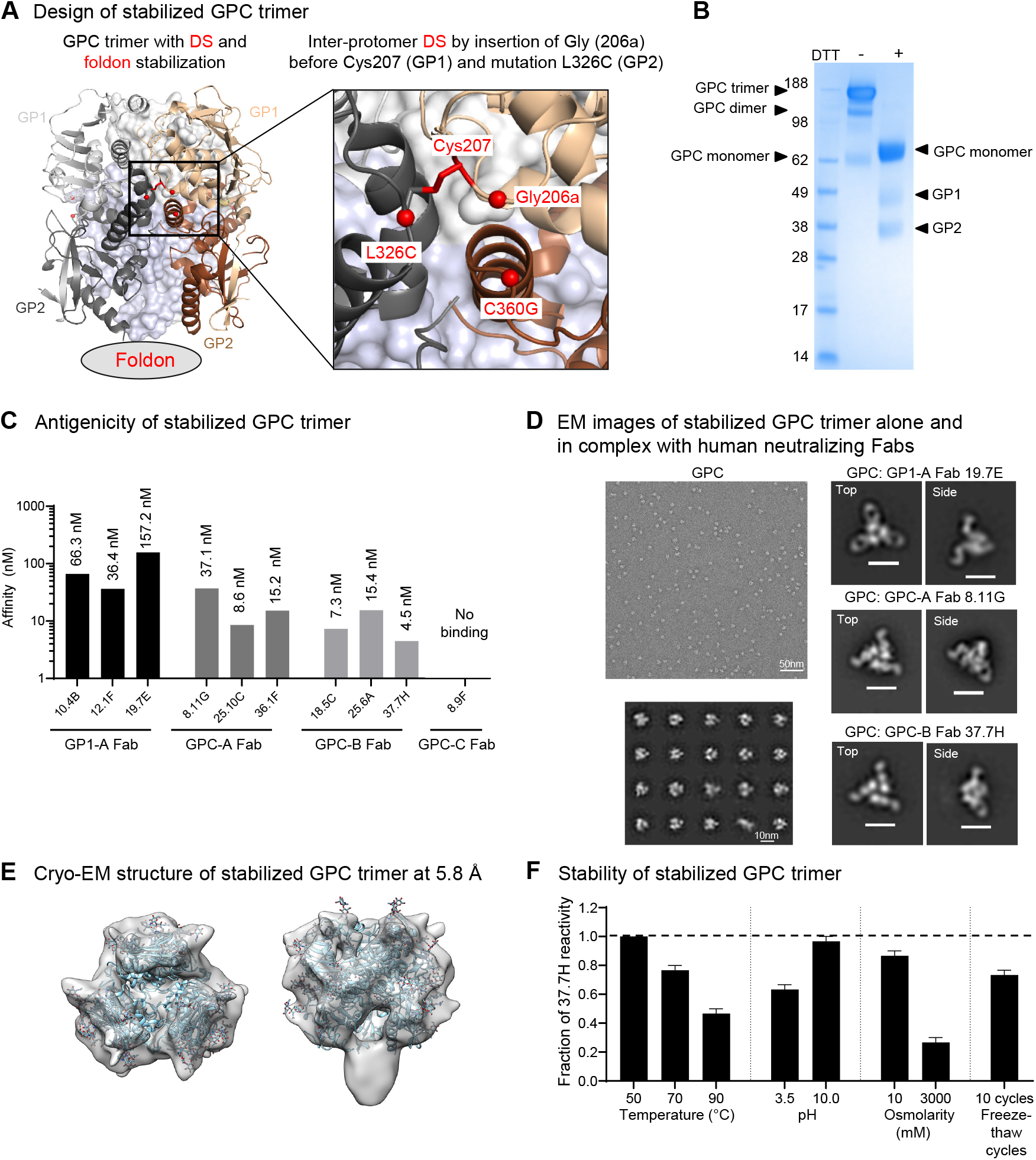
Design and characterization of stabilized soluble Lassa GPC trimer. (A) Structure-based design of stabilized soluble Lassa GPC trimer. An inter-protomer disulfide (DS) bond linked GP1 of one protomer to GP2 of a neighboring protomer and a foldon domain was appended to the C-terminus of GP2. Inset shows a view of the inter-protomer DS shown as spheres. The two protomers are shown as ribbons in light gray (GP1) and dark gray (GP2), and wheat (GP1) and brown (GP2). (B) SDS-PAGE of stabilized Lassa trimer under non-reducing and reducing conditions. A high molecular weight band three times of the monomeric form was observed in the non-reducing condition. (C) Binding affinity of the stabilized Lassa GPC trimer toward Fabs of four groups of human Lassa-neutralizing antibodies, GP1-A, GPC-A, GPC-B, and GPC-C. (D) Negative-stain EM images of the stabilized Lassa GPC trimer alone and in complex with human neutralizing Fabs. Representative top view and side view are shown. (E) Cryo-EM structure of the stabilized Lassa GPC at 5.8 Å confirmed the trimeric association of the protein. (F) Physical properties of the stabilized Lassa GPC trimer. Stability of the stabilized trimer was assessed as fractional binding reactivity to 37.7H after treatments under various temperature, pH, osmolarity changes and freeze-thaw cycles. Triplicate measurements were made, and results were represented as mean ± SEM. The dotted line shows the antibody reactivity of trimer prior to physical stress.

The resultant LASV GPC trimer expressed as a soluble protein with a final yield of approximately 0.5 mg/L by transient transfection of mammalian cells **(Fig. S2)**. The purified protein yielded a major band at an expected size of a trimer (∼200 kDa) on SDS-PAGE in the absence of reducing agent, indicative of the formation of inter-protomer disulfide bond **(Fig. 1B)**. In the presence of reducing agent, bands for GP1 and GP2 appeared; however, the majority of the stabilized trimer existed as a band of the size for a protomer, indicating that GP1 and GP2 subunits were not efficiently cleaved **(Fig. 1B)**. To verify the prefusion conformation of the stabilized LASV GPC trimer, we analyzed its antigenicity by bio-layer interferometry (BLI) assay for recognition by a panel of 10 human LASV neutralizing antibodies from four epitope groups; all except for the GPC-C antibody 8.9F could bind, suggesting that the stabilized trimer possessed generally similar antigenic property as the previously published GPCysR4 construct (8) **(Fig. 1C)**. Negative-stain electron microscopy (EM) confirmed the homogeneous size and shape expected for a stable trimer in prefusion conformation of this stabilized GPC **(Fig. 1D)**. More importantly, in the presence of various human neutralizing Fabs, the stabilized trimer preserved its trimeric shape **(Fig. 1D)**. To further examine the overall architecture of the inter-protomer disulfide-stabilized LASV GPC trimer, we determined a cryo-EM structure at 5.8 Å resolution from 47,597 particles **(Fig. 1F)**. We observed trimers with C3 symmetry but the majority of the particles displayed C1 symmetry. Lastly, the stabilized GPC trimer could withstand physical stresses under various temperature (50°C–90°C), pH (3.5 and 10), osmolarity (10 mM and 3000 mM NaCl), and freeze-thaw conditions, as evidenced by the retained 37.7H reactivity (8) after treatments **(Fig. 1D)**. Overall, our design yielded a soluble prefusion-stabilized LASV GPC, which exhibited desired antigenic and structural characteristics.

### Identification of LASV GPC-binding nanobodies from camel and shark libraries

To identify single domain antibodies capable of binding our prefusion-stabilized LASV GPC trimer, we screened phage display libraries of variable domain of New Antigen Receptors (V_NAR_) from sharks (21, 22) and single variable domain on heavy chain (V_H_H) antibodies from camels (24). After four consecutive rounds of panning, phage was enriched by 400-1200-folds for binding the stabilized LASV GPC trimer **(Fig. S3)**. At the end of the fourth round of panning, we identified four individual clones (B8, B10, C3 from shark V_NAR_, and D5 from camel V_H_H that exhibited enhanced binding to stabilized LASV GPC trimer in ELISA, whereas no binding was observed with the control protein bovine serum albumin **(Fig. 2A)**. The binding affinities of these single domain antibodies to the stabilized GPC trimer were in the range of 19-44 nM as measured by BLI **(Fig. 2B)**. Binding competition revealed that C3 and D5 competed for binding to GPC with known human LASV neutralizing antibodies from the GPC-B group but not from groups GP1-A and GPC-A B8, and B10 showed only mild competition with the human neutralizing antibodies for binding the GPC trimer **(Fig. 2C)**.

**Fig. 2.**
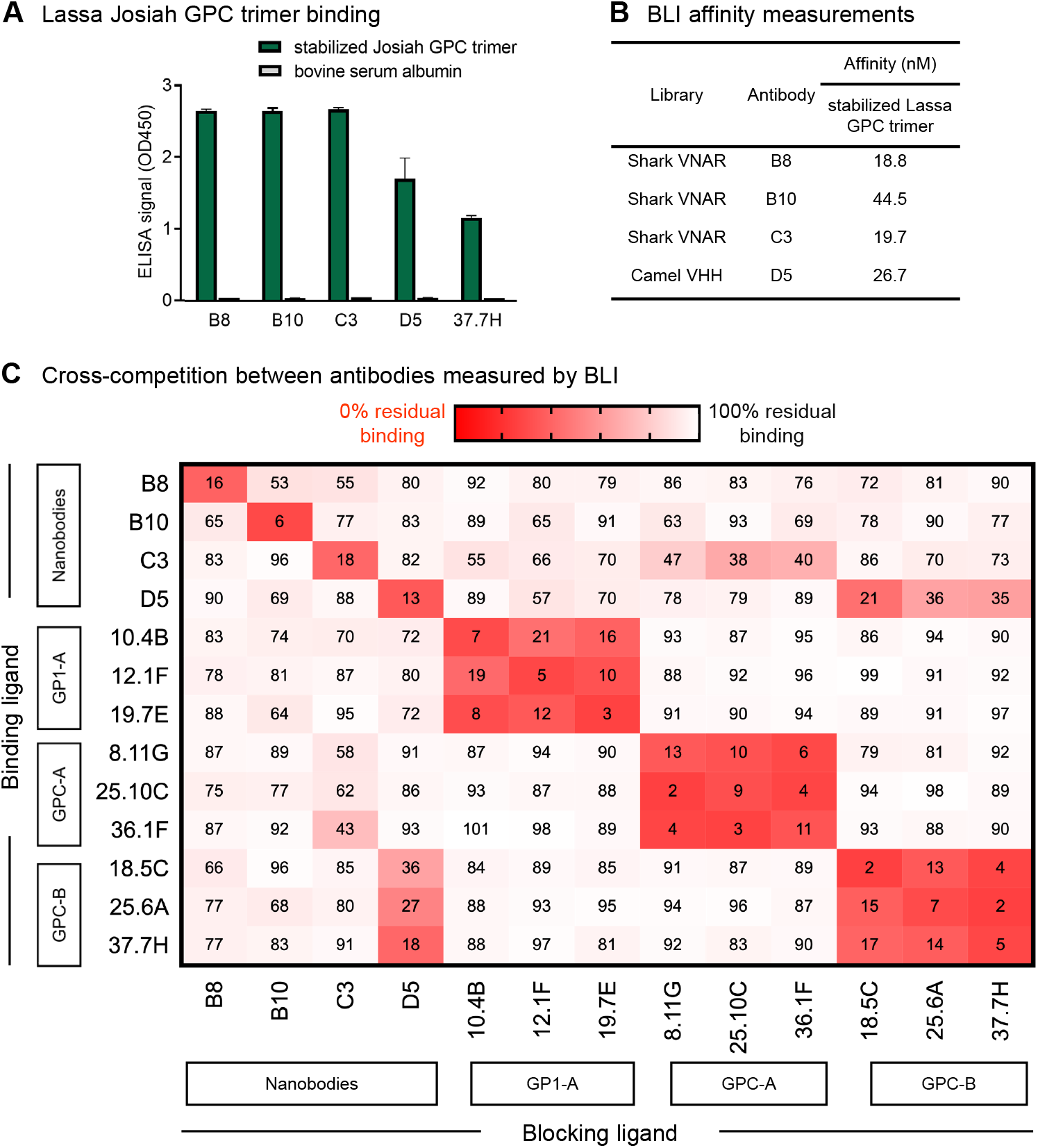
Single domain antibodies identified from camel and shark library panning bind prefusion-stabilized Lassa GPC. (A) Three single domain antibodies from shark (B8, B10, C3) and one from camel (D5) libraries showed binding to the stabilized Lassa GPC trimer by ELISA. A potent human Lassa neutralizing antibody, 37.7H, was used as a positive control. Select nanobodies showed minimal reactivity toward bovine serum albumin (BSA). Triplicate measurements were made and results were represented as mean + SEM. (B) Binding affinities of the six single domain antibodies toward the stabilized GPC trimer. (C) Cross-competition between the four single domain antibodies and human LASV-neutralizing antibodies toward the stabilized Lassa GPC trimer. Epitope binning was performed using biolayer interferometry. His-tagged stabilized Lassa GPC trimer was loaded onto the NTA sensor tips; then the blocking ligand was loaded, followed by loading of the second ligand. The numerical data indicate percent binding of the binding ligand in the presence of the blocking ligand.

### Superior LASV neutralization by nanobody-IgG2a and most human neutralizing antibodies, except the quaternary-specific GPC-B group, attributed to avidity

We next assessed the single domain antibodies for neutralizing activity against pseudotyped LASV Josiah strain. At the tested concentration range (1 µg/ml – 1 mg/ml), all four single domain antibodies failed to show >50% neutralization **(Fig. 3A)**. Since improved neutralization potency with multimeric single domain antibodies have been reported for other viruses (25-28), we linked a single domain antibody molecule to a llama IgG2a hinge region and then to the Fc domain of human IgG1 (29, 30). We found that three of the four bivalent IgG2a antibodies exhibited modest neutralization toward the Josiah pseudovirus with IC_50_ ranging from 12 to 260 µg/ml **(Fig. 3B)**. Of these bivalent IgG2a antibodies, D5-IgG2a demonstrated the best neutralizing activity with an IC_50_ of 12 µg/ml **(Fig. 3B)**.

**Fig. 3.**
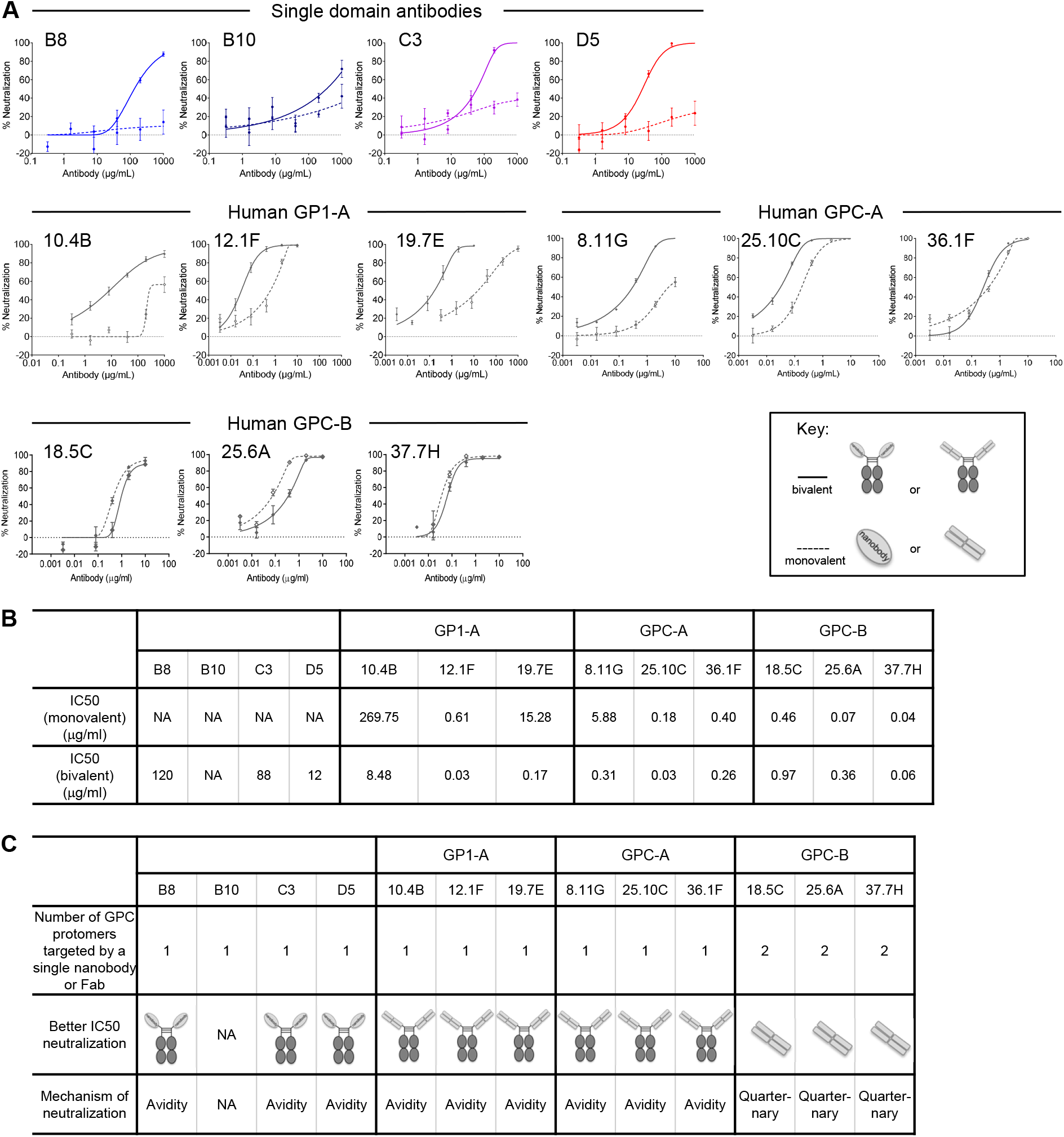
Neutralization of Lassa virus by single domain antibodies and most human neutralizing antibodies requires avidity. (A) Neutralization of pseudotyped Josiah strain of Lassa virus by single domain antibodies and human Lassa neutralizing antibodies in both monovalent (dotted line) and bivalent (solid line) formats. (B) Summary of the IC50 values of single domain antibodies and human Lassa neutralizing antibodies in monovalent and bivalent formats. (C) Summary of the proposed neutralization mechanisms of single domain antibodies and human antibodies for Lassa virus.

To delineate the molecular mechanism for LASV neutralization by nanobody-IgG2a, we compared the neutralization potencies of D5 and the human neutralizing antibodies in monovalent and bivalent context against the pseudotyped Josiah virus. As described above, D5 could neutralize LASV only in a bivalent IgG2a format **(Fig. 3A, top left)**. Such a trend was similar to the human GP1-A and GPC-A groups of antibodies, for which higher neutralization potency was observed in bivalent IgGs than in monovalent Fabs **(Fig. 3A, top right and lower left)**, but was opposite to the human GPC-B group of antibodies, for which better neutralization abilities were observed in monovalent Fabs than bivalent IgGs **(Fig. 3A, lower right)**. Although the neutralization potency of D5-IgG2a was weaker than the human neutralizing antibodies **(Fig. 3B)**, this demonstrates the feasibility of using prefusion-stabilized GPC to identify non-human neutralizing antibodies *in vitro*. Overall, the results indicated that D5 in bivalent IgG2a format, similar to human GP1-A and GPC-A antibodies, neutralized LASV via avidity (31, 32), a mechanism different from that employed by the most common group of LASV GPC-B antibodies, where a single Fab was able to bind and engage two protomers in a quaternary manner **(Fig. 3C)**.

### Cryo-EM structure of LASV GPC in complex with nanobody D5 and 8.11G Fab

We sought to visualize binding by the most potent of the nanobodies, D5. We were unable to interpret cryo-EM ima]ges of D5 bound to GPC, but the addition of human GPC-B antibody 8.11G lead to better resolved reconstructions, enabling determination of the cryo-EM structure at 4.7 Å of our prefusion-stabilized LASV GPC trimer in complex with a single D5 nanobody and two GPC-A 8.11G Fabs **(Figs. 4A and S4, Table 1)**. D5 bound to the apex of the GPC trimer, forming asymmetric interactions with all three GPC protomers **(Fig. 4B)**. 8.11G bound to an interface between GP1 and GP2 subunits of single protomer **(Fig. 4C)**. Despite the heavy glycosylation of GPC 8.11G navigated through the shield, contacting six glycans.

**Table 1.**
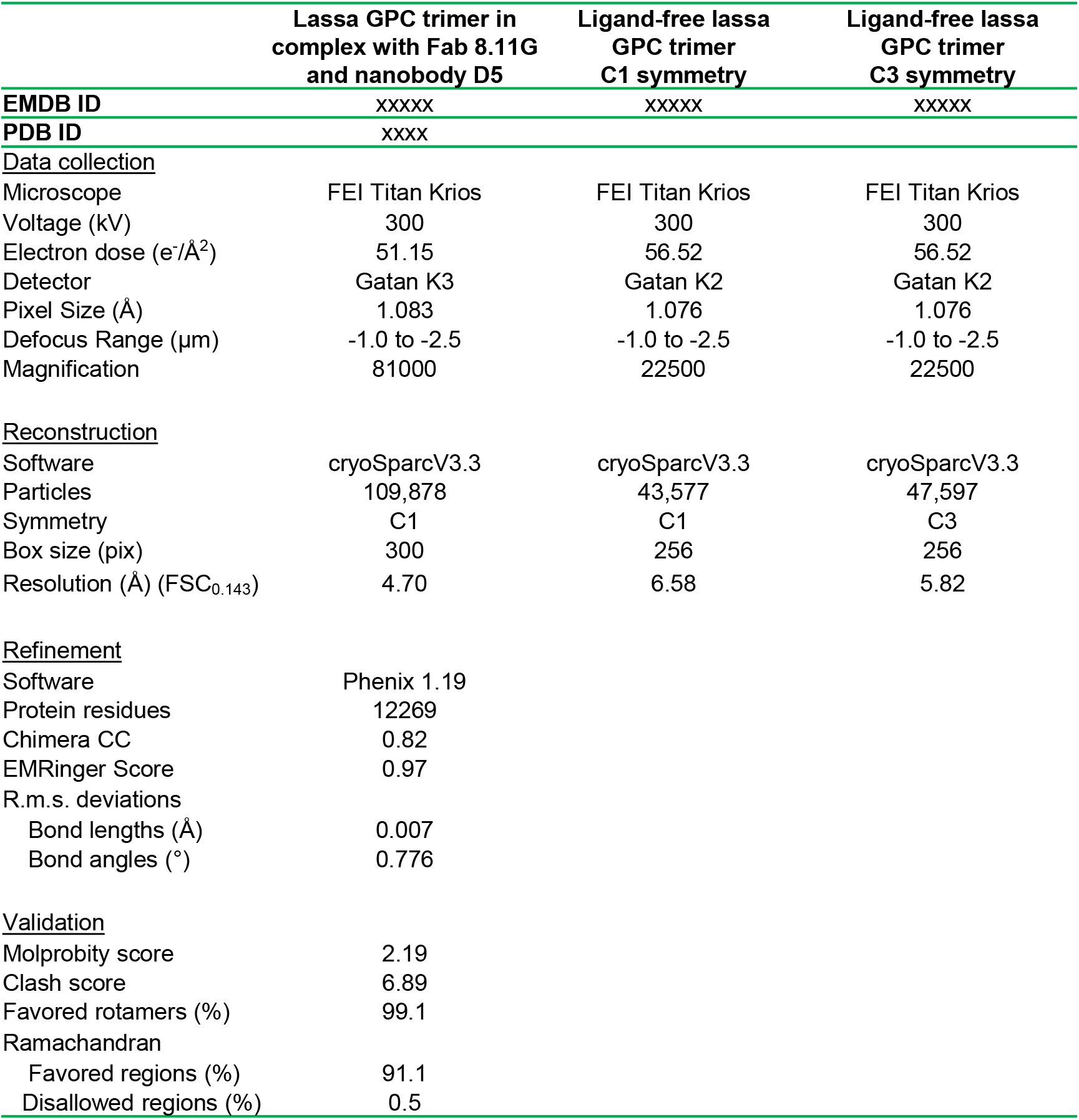
Cryo-EM data, reconstruction, refinement, and validation statistics.

**Fig. 4.**
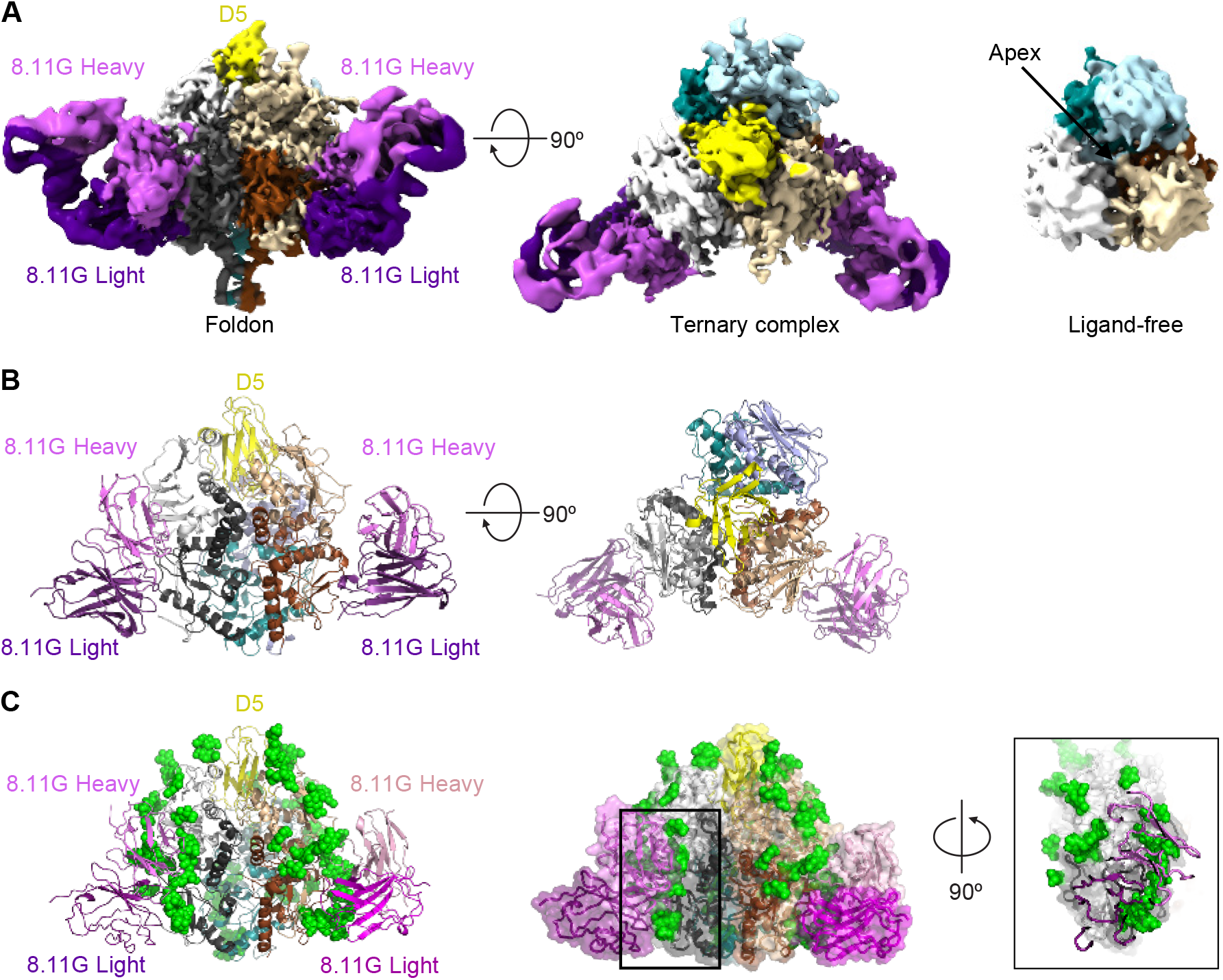
Cryo-EM structure of D5 with Lassa trimer reveals details of 8.11G and D5 recognition. (A) CryoEM density is shown for a complex of the stabilized GPC trimer bound to two Fabs of 8.11G and a single D5. An unliganded trimer is shown in the right panel highlighting the cavity where D5 binds. (B) The atomic model is shown in cartoon representation. A 5Å footprint of D5 is shown, highlighting interactions with all three protomers. (C) The highly glycosylated epitope of 8.11G is highlighted. Glycans for which density was observed are displayed as green spheres. The structure is displayed in cartoon format (left) and with a transparent surface (middle). (right) A view along the axis of the 8.11G Fab binding with Fab shown as cartoon and the GPC protein and glycans shown in surface representation.

The structure of the trimer bound by D5 displayed an asymmetric assembly and when compared to the crystallized GPCs (8, 20), a single protomer aligns with an rmsd of 1.5Å after removing outliers, however the neighboring protomers extended over 17 Å farther at certain regions near the apex with overall rmsds of 13.2 Å and 21.9 Å **(Fig. 5A)**. This extension resulted in the allosteric disruption 37.7H binding, and stabilization of this conformation by D5 would prevent 37.7H from binding, thereby providing an explanation for the observed competition between D5 and GPC-B antibodies, which bind across adjacent GPC protomers distal from the apex **(Fig. 5B)**. The asymmetry in the protomer interface extensions was matched by the protrusion of an uncleaved furin site, between GP1 and GP2 **(Fig. 5C)**. This results in two internal sites and a single protruding loop, disrupting the trimer symmetry and inflating the radius of the apex chalice targeted by D5.

**Fig. 5.**
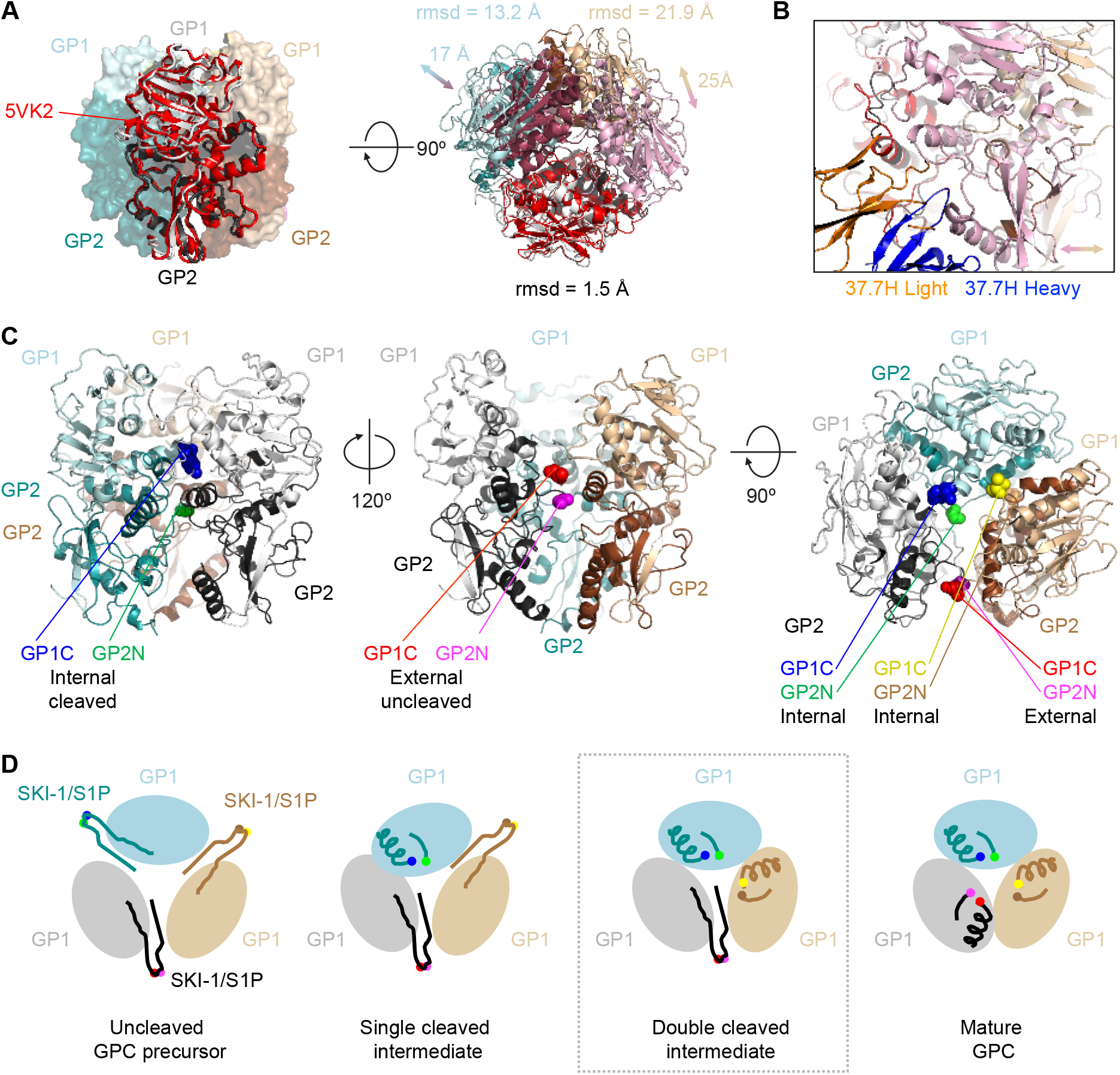
Analysis of asymmetric Lasa GPC trimer reveals a cleavage-intermediate site of vulnerability at the trimer apex. (A) A single protomer of the GPC trimer shows a closely matching RMSD with that of the C3 symmetric GPC (PDB ID 5VK2, colored red, pink, and raspberry). The C1 symmetric GPC trimer observed here does not maintain the same quaternary assembly with the adjacent protomer oriented to accommodate the uncleaved peptide. After removing outlying residues the rmsd for one GP1/GP2 across 294 Cα atoms of the protomer is 1.5 Å, however the rmsd’s for the other two protomers are 13.2 and 21.9 Å across 296 and 294 Cα, respectively. Specific apex regions showing distances from 17-25 Å. (B) Explanation for the loss of one of the three 37.7H binding sites on the trimer with D5 stabilizing a more open interface between the protomers. (C) The Lassa GPC is shown with focus on one of the two cleaved protomers (left) with the internal termini of GP1 (light cyan) and GP2 (teal) shown. Rotation by 120º shows the external location of the uncleaved peptide. (D) A schematic representation of the cleavage intermediates in the maturation of the GPC trimer is shown from a top view looking down the trimer axis. The SKI-1/S1P cleavage site must be cleaved on all three protomers to enable a tightly packed GPC trimer. The neutralizing nanobody D5 binds the first three populations (uncleaved, single-cleavage and double-cleaved).

### An apex-situated site of vulnerability centers on the receptor binding site

We evaluated the D5-defined site of vulnerability at the GPC trimer apex for its potential utility as a vaccine target. Residues within a 5Å footprint of the nanobody were encompassed in all three protomers with buried surface areas of 730, 620, and 350 Å^2^ (**Fig. 6A**). The asymmetrically opened conformation, defined by the binding of D5, revealed an apex site with little steric hindrance from the apex glycans (**Fig. 6B**).

**Fig. 6.**
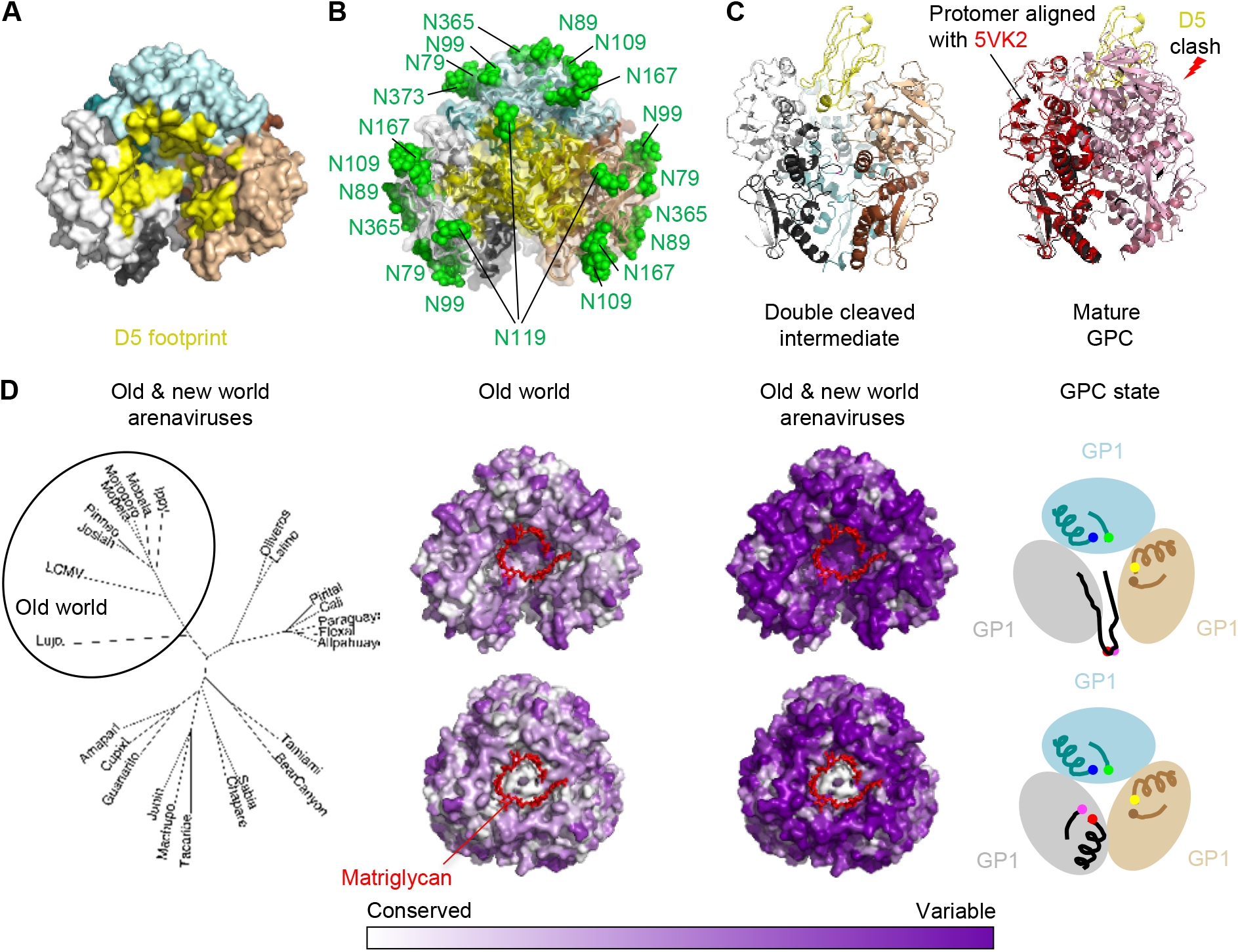
Apex site of nanobody vulnerability encompasses binding site for matriglycan receptor. (A) A surface representation of the cleavage intermediate trimer is shown. Residues within a 5Å footprint of the D5 nanobody are highlighted in yellow. (B) A transparent surface representation is shown as in panel A with cartoon representation underneath. Glycans are displayed as green spheres. (C) D5 fits into pocket at apex of double-cleaved intermediate, but not into the mature GPC, in which one protomer orientation shifts the apex region by >10 Å (pink) and the second protomer shifts at apex regions by >20 Å (raspberry), sterically blocking access to the the D5 binding site. (D) (left) A phylogenetic tree of select arenavirus GPC sequences is displayed to illustrate the overall sequence diversity and highlight the clustering of old world versus new world virus. (right) Conservation of the GPC residues is shown for the Old world viruses versus all arenaviruses with the double-cleavage intermediate on the top and the mature GPC on the bottom, the matriglycan (red) from PDB ID 7PVD is shown on all surfaces.

When compared to the mature closed structures (8, 12, 20), D5 would be obstructed from binding at the apex (**Fig. 6C**). Notably, this cavity also serves as the receptor binding site. Sequence analysis of representative New and Old world arenaviruses displayed an expected clustering and conservation of these sequences was mapped onto the surface of the GPC structures (**Fig. 6D**). Greater conservation was displayed in the GP2 subunit than the GP1, even those residues around the binding site except for the internal cleavage residues at the center of the apex cavity, which make pivotal contacts with the matriglycan receptor (**Fig. 6D**). The revealed site was partially conserved, especially with Old world arenaviruses, but not arenaviruses in general (**Fig. 6D**), where the major source of conservation involved the conserved cleavage site between gp1 and gp2. Interestingly, the apex site was less conserved in the double-cleaved intermediate, as the cleavage sites are not at the apex; overall, the apex site was conserved within all Lassa virus lineages, but not among more diverse arenaviruses.

## DISCUSSION

The LASV GPC trimer is metastable, conformationally labile, and heavily glycosylated, rendering the elicitation of neutralizing antibodies difficult (19). Stabilization by human neutralizing antibodies of the GPC-B class have enabled the structural analysis of the LASV GPC trimer (8, 20), yet a stabilized stand-alone GPC trimer has not been generated for use as immunogen. Here, we employed structure-based design strategy to engineer an inter-protomer disulfide bond and a foldon trimerization domain to obtain a soluble ligand-free LASV GPC trimer stabilized in its prefusion state, which demonstrated similar antigenicity and trimeric architecture as the previously published antibody-bound LASV GPC structure. Panning of this stabilized LASV GPC trimer against phage libraries identified a single domain antibody, D5, which bound the stabilized trimer and neutralized LASV in a bivalent IgG2a format, recognizing a glycan-free hole at the trimer apex. The apex-situated site of vulnerability appeared to be specific to GPCs with partial cleavage between GP1 and GP2 subunit. These results demonstrate the utility of the stabilized LASV GPC trimer for identification of LASV-neutralizing antibodies and reveal an apex-situated site of vulnerability.

Other potently neutralizing antibodies have been identified, which – like D5 - bind with a one-antibody-per trimer stoichiometry at the trimer apex and neutralize potently. These include the very potent PG9 (33), PGT145 (34), and CAP256-VRC25.26 (35, 36), which neutralize HIV-1 potently, as well as the potent antibodies PIA174 (37) and PI3-E12 (38), which neutralizes PIV3 potently. The mechanism of such antibodies offers a three-fold advantage in stoichiometry, and therefore, potentially, effective concentration necessary for neutralization. Such antibodies, rather than occluding receptor binding, are able to prevent conformational triggering to fusion competent forms.

The cryo-EM structure of D5 bound to the GPC trimer highlighted an asymmetric cleavage intermediate that exposed a new site of vulnerability at the trimer apex. We speculate that this conformation is available in several of the intermediate stages during the SP1 cleavage process. It is clear, however, that this conformation must be present on GPC trimers on the virus surface since D5 is able to neutralize virus in the IgG2a format. Native trimer instability may play a role in exposing this vulnerable conformation on the virion surface. Overall, we have stabilized a GPC trimer antigen in the absence of antibody, screened for a potently neutralizing nanobody, and determined an asymmetric trimer structure with a novel apex site of vulnerability that could be developed as an immunogenic target for a Lassa virus vaccine.

## Materials and Methods

Designs of inter-protomer disulfide bond and foldon trimerization domain in LASV GPC were performed based on the LASV GPC-37.7H Fab complex structure (5VK2). Proteins were expressed in 293F cells transfected with Turbo293 transfection reagent (SPEED BioSystem) and then purified by a serial His6-strepTagII purification approach. Purified recombinant LASV GPC protein was analyzed antigenically by Octet interferometry with a panel of human neutralizing antibodies, and structurally by negative-stain EM. Cryo-EM data for the stabilized LASV GPC trimer protein were collected using a Titan/Krios microscope with Leginon and map reconstruction carried out by cryoSPARC. The relative binding epitopes of single domain antibodies isolated from phage display library panning were mapped by cross-competition assay with the human LASV neutralizing antibodies. Neutralizing activities of bivalent single domain antibodies and human LASV neutralizing antibodies were assessed by pseudotyped LASV neutralization assay. Details are provided in *SI Materials and Methods*.

## ACKNOWLEDGEMENTS

We thank M. Sastry for assistance with control antibodies, J. Stuckey for assistance with figures, and members of the Structural Biology Section and Structural Bioinformatics Core, Vaccine Research Center, for discussions and comments on the manuscript. We also thank members of the Electron Microscopy Group at the New York Structural Biology Center for assistance with data collection. Support for this work was provided by the Intramural Research Program of the Vaccine Research Center, National Institute of Allergy and Infectious Diseases (NIAID), National Institutes of Health (NIH) and by the Intramural Research Program of the Center for Cancer Research, National Cancer Institute, NIH. This work was also supported in part with federal funds from the Frederick National Laboratory for Cancer Research, NIH, under contract HHSN261200800001E. Some of this work was performed at the Simons Electron Microscopy Center and National Resource for Automated Molecular Microscopy located at the New York Structural Biology Center, supported by grants from the Simons Foundation (SF349247), NYSTAR, and the NIH National Institute of General Medical Sciences (GM103310).

