## Supporting Information for "Prefusion-Stabilized Lassa Virus Trimer Identifies Neutralizing Nanobodies and Reveals an Apex-Situated Site of Vulnerability"

**SI Materials and Methods:**

**Structure-based design of stabilized LASV GPC trimer**. 164 design variants based on the published LASV GPC-37.7H complex crystal structure (PDB ID 5VK2) (1) were made. In total, we designed 68 disulfide bonds, 43 cavity-filling mutations, 22 helix-breaking mutations, and 31 trimerization domain insertions with various length linkers (Fig. S1).

**Antigenic screening of LASV GPC stabilizing designs**. Assessment of all 164 constructs was perfoIrmed using high-throughput 96-well microplate expression format followed by an ELISA-based antigenic evaluation as described previously (2). Briefly, $2.5x10^5$ cells/ml of HEK 293T cells (Thermo Fisher Scientific, MA) were seeded in a 96-well microplate and cultured in expression medium (high glucose DMEM supplemented with 10% ultra-low IgG fetal bovine serum and 1x-non-essential amino acids) at 37°C, 5% $CO_2$ for 20 h. Plasmid DNA and Turbo293 transfection reagent (Speed BioSystems) (3) were mixed and added to the cells. One day post transfection, enriched medium (high glucose DMEM plus 25% ultra-low IgG fetal bovine serum, 2x nonessential amino acids, 1x glutamine) was added to all wells. The cells were cultured at 37°C, 5% $CO_2$ for five days. Supernatants with the expressed LASV GPC variants were harvested and tested by ELISA for binding to 37.7H antibody using $Ni^{2+}$-NTA microplates.

**Enzyme-linked immunosorbent assay ELISA**. $Ni^{2+}$-NTA microplates (Pierce) were coated with 100 µl/well the supernatant of LASV GPC variants at 4°C overnight. After standard washing procedures with 0.1% Tween 20 in PBS, 100 µl of 10 µg/ml GPC-specific antibody 37.7H was added to the wells and incubated at room temperature for 2 h. After that, 100 µl of horseradish peroxidase (HRP)-conjugated goat anti-human IgG Fc antibody (1:5000 v/v)

(Jackson ImmunoResearch Laboratories Inc) was introduced to the plate for incubation at room temperature for 1 h. Subsequently, plates were washed and then developed with 3,3´,5,5´-tetramethylbenzidine (TMB) substrate (Kirkegaard & Perry Laboratories). Signal was read at 450 nm by a plate reader (Beckman Coulter).

**LASV GPC trimer protein expression and purification**. Disulfide and foldon-stabilized LASV GPC sequence was attached to a thrombin cleavage sequence, a hexahistidine tag, and a Strep-tag at its C-terminal end. The stabilized LASV GPC protein was expressed by transient transfection in 293F cells (Thermo Fisher) with Turbo293 transfection reagent (SPEED BioSystem) using the established protocol (4). Briefly, one liter of 293F cells at a density of 1.2 $\times 10^6$ cells/ml were co-transfected with 700 μg/liter of the LASV GPC expression plasmid and 300 μg/liter of furin plasmid. 6 days post transfection, the culture supernatant was harvested and protein was purified from the supernatants by Nickel- (Roche) and Strep-Tactin- (IBA lifesciences) affinity columns. The resulted protein was loaded on a Superdex 200 16/600 size exclusion column (GE Healthcare) to be further polished for use in subsequent assays.

**Production of human LASV antibodies**. Immunoglobulin heavy chain or light chain sequences were constructed by gene synthesis and then cloned into human IgG1, lambda, or kappa expression plasmids as previously described (5). Heavy and light chain expression plasmid DNA was transfected into Expi293F cells (Thermo Fisher) in 1:1 (v/v) ratio using Turbo293 transfection reagent (3). Monoclonal antibodies from the culture supernatants were purified using recombinant Protein-A Sepharose (GE Healthcare) as per the manufacturer’s instructions.

**Antibody Fab preparation**. The purified human IgG proteins were cleaved by LysC enzyme (1:4000 w/w) (Roche) at 37$^{o}$C overnight to yield Fabs. On the next day, the enzymatic digestion reaction was terminated by addition of protease inhibitor (Roche). The cleavage mixture was then passed through a Protein-A column to separate the Fc fragments from the Fab. The Fab collected in the flow-through was loaded onto a Superdex 200 16/60 column for further purification.

**LASV GPC antigenic characterization**. An Octet Red384 instrument (fortéBio) was used to measure the binding kinetics between the stabilized LASV GPC trimers and human LASV neutralizing antibodies or nanobodies. Assays were performed at 30°C in tilted black 384-well plates (Geiger Bio-One). Ni-NTA sensor tips (fortéBio) were used to capture the histidine-tagged stabilized LASV GPC trimer for 300 s. Then, the biosensor tips were equilibrated for 60 s in PBS before measurement of association with antigen-binding fragments (Fabs) in solution (6.25 nM to 400 nM) for 180 s. Subsequently, Fabs were allowed to dissociate for 300s. Parallel correction to subtract systematic baseline drift was carried out by subtraction of the measurements recorded for a loaded sensor dipped in PBS. Data analysis and curve fitting were carried out using the Octet Data Analysis Software 9.0 (fortéBio). Experimental data were fitted with the binding equations describing a 1:1 interaction. Global analysis of the data sets assuming reversible binding (full dissociation) were carried out using nonlinear least-squares fitting allowing a single set of binding parameters to be obtained simultaneously for all of the concentrations used in each experiment.

**Negative-stain electron microscopy**. The protein was diluted with a buffer containing 10 mM HEPES, pH 7.0, and 150 mM NaCl to a concentration of 0.02 mg/ml and adsorbed to a freshly glow-discharged carbon-coated copper grid. The grid was washed with the same buffer, and the adsorbed protein molecules were negatively stained with 0.7% uranyl formate. Micrographs were collected at a nominal magnification of 100,000 using SerialEM (6) on an FEI T20 electron microscope equipped with a 2k x 2k Eagle camera and operated at 200 kV. The pixel size was 0.22 nm. Particles were picked automatically using in-house written software (YT, unpublished) and extracted into 100x100-pixel boxes. Reference-free 2D classifications were performed using Relion (7).

**Physical stability of the designed LASV GPC trimer**. To assess the physical stability of the designed LASV GPC trimer under various stress conditions, we treated the proteins with a variety of pharmaceutically relevant stresses such as extreme pH, high temperature, low and high osmolarity, and repeated freeze/thaw cycles while at a concentration of 50 μg/ml. The physical stability of treated LASV GPC trimer was evaluated by the preservation of binding to the GPC-specific antibody 37.7H. Temperature treatments were carried out by incubating the stabilized LASV GPC protein solutions at 50°C, 70°C and 90°C for 60 m in a PCR cycler with heated lid. In pH treatments, the stabilized LASV GPC protein solution was adjusted to pH 3.5 and pH 10.0 with appropriate buffers for incubation at room temperature for 60 m and subsequently neutralized to pH 7.5. In osmolarity treatments, the stabilized LASV GPC protein solutions originally containing 150 mM NaCl were either diluted with 2.5 mM Tris buffer (pH 7.5) to an osmolarity of 10 mM NaCl or adjusted with 4.5 M $MgCl_2$ to a final concentration of 3.0 M $MgCl_2$. Protein solutions were incubated for 60 minutes at room temperature and then returned to

150 mM salt by adding 5.0 M NaCl or dilution with 2.5 mM Tris buffer, respectively, and concentrated to 50 μg/ml. The freeze/thaw treatment was carried out by repeatedly freezing the stabilized LASV GPC protein solutions in liquid nitrogen and thawing at 37°C ten times. The degree of physical stability is reported as the ratio of steady state 37.7H antibody-binding level before and after stress treatment.

**Phage display panning of nanobody libraries**. Shark $V_{NAR}$ and camel $V_HH$ nanobody phage display libraries were constructed in Dr. Mitchell Ho’s laboratory at the National Cancer Institute (Bethesda, Maryland) according to the protocol described previously (8). The phage panning protocol has been described previously (9, 10). Briefly, an immunotube (Nunc/Thermo Fisher Scientific, Rochester, NY) was coated with 0.5 ml of 10 μg/ml LASV GPC trimer in PBS at 4°C overnight. After decanting the coating buffer, the immunotube was treated with 0.5 ml blocking buffer (10% milk in PBS) at room temperature for 1 h. Then, a fixed amount of input phage from the shark or camel libraries was added to the immunotube for binding to the LASV GPC trimer at room temperature for 2 h with gentle shaking. The immunotube was washed with PBS containing 0.05% Tween-20 to remove unbound phages. Subsequently, the bound phages were eluted with 100 mM triethylamine. At each round of panning, output phage enrichments were assessed and monitored by polyclonal phage ELISA. Single colonies were picked at the final round of panning for DNA sequencing. The binding ability of the identified nanobodies from phage display toward the stabilized LASV GPC trimer was further evaluated by ELISA, with bovine serum albumin (BSA) serving a negative control.

**Polyclonal phage ELISA**. A 96-well plate (Corning) was coated with 50 µl/well of 5 µg/ml stabilized LASV GPC trimer in PBS buffer at 4°C overnight. After blocking with 3% milk in 100% superblock buffer (Thermo Scientific) at room temperature for 2 h, 50 µl phage were added to the plate and incubated at room temperature for another hour. Binding of phage to the stabilized LASV GPC was detected by HRP-conjugated anti-M13 antibody (GE Healthcare). The cut-off value for positive binder was set as 3× higher signal of antigen binding compared to background signal.

**Expression of nanobodies**. The pComb3x phagemids containing the $V_{NAR}$ or the $V_HH$ binders were transformed into HB2151 *E. coli* cells. The formed colonies were pooled for culture in 2 L 2YT media containing 2% glucose, 100 µg/ml ampicillin at 37° C until the OD600 reaches 0.8–1. Culture media was then replaced with 2YT media containing 1 mM IPTG (Sigma), 100 µg/ml ampicillin, and shook at 30°C overnight for soluble protein production. Bacteria pellet was spun down and lysed with polymyxin B (Sigma Aldrich) for 1 h at 37° C to release the soluble protein. The supernatant was harvested after lysis and purified using HisTrap column (GE Healthcare) using AKTA.

**Protein ELISA**. Protein ELISA was used to evaluate the binding ability of the selected nanobody binders toward the stabilized LASV GPC trimer. Briefly, a 96-well plate was coated with either the stabilized LASV GPC trimer or BSA at 5 µg/ml in PBS, 50 µl/well, at 4°C overnight. After blocking with 100% superblock buffer, the nanobodies were diluted into 1 µg/ml using 10% PBST in 100% superblock and then added to the plate for incubation at room temperature for 1 h. Binding signal was detected by HRP-conjugated anti-Flag antibody (Sigma).

**Cross-competition assay**. The histidine- and Strep-tagged stabilized LASV GPC trimer protein (30 μg/ml) was captured by a mouse anti-streptavidin antibody, which was immobilized by the anti-mouse Fc sensor tips to a final mean signal level of 1.0-1.5 nm. The trimer-coated tips were then dipped into either PBS or the pre-determined saturating concentrations of Fabs (1000 nM) or nanobodies (500 nM) (first ligand) in PBS for 300 s. After loading, the sensor tips were incubated in PBS briefly for 60 s to remove unbound ligands for baseline adjustment. Subsequently, the sensor tips were dipped into wells containing a fixed concentration of competing ligands (second ligand, 1000 nM Fabs or 500 nM nanobodies) for another 300 s, followed by 300 s of dissociation in PBS. Raw data was processed using Octet Data Analysis Software 9.0. Percent of residual binding was calculated as follows: (response signal from the second ligand in presence of first ligand / response signal from the second ligand in absence of first ligand) x 100.

**Generation of nanobody-IgG2a proteins**. To express nanobodies in a bivalent IgG format, the gene encoding nanobody variable region was cloned into the mammalian protein expression vector pVRC8400 in front of DNA sequences encoding an Ala-Ala-Ala linker, the llama IgG2a hinge sequence (EPKIPQPQPKPQPQPQPQPKPQPKPEPECTCPKCP) and the human IgG1 Fc domain. The nanobody IgG2a proteins were expressed by transient transfection in 293F cells (Thermo Fisher) with Turbo293 transfection reagent (SPEED BioSystem) using the protocol described above and purified by protein A affinity column.

**Production of pseudovirus**. Recombinant Indiana VSV (rVSV) expressing LASV GPC were generated as previously described (12, 13) . HEK293T cells were grown to 80% confluency before transfection with plasmids expressing LASV Josiah GPC using FuGENE 6 (Promega). Cells were cultured at 37°C with 5% $CO_2$ overnight. The next day, medium was removed and VSV-G pseudotyped ΔG-luciferase (G*ΔG-luciferase, Kerafast) was used to infect the cells in DMEM at a MOI of 3 for 1 h before washing the cells with 1× DPBS three times. DMEM supplemented with 2% fetal bovine serum and 100 I.U./mL penicillin and 100 µg/mL streptomycin was added to the infected cells and they were cultured overnight as described above. On the following day, the supernatant was harvested and clarified by centrifugation at 300g for 10 m before aliquots and storage at -80° C.

**Pseudovirus-based neutralization assay**. Neutralization assays were performed by incubating pseudoviruses with serial dilutions of antibodies and measured by the reduction in luciferase gene expression. In brief, Vero E6 cells (ATCC) were seeded in a 96-well plate at a concentration of $2 \times 10^4$ cells/well the day before. Pseudoviruses were incubated with serial dilutions of antibodies (six dilutions in a 5-fold stepwise manner) in triplicate at 37° C for 30 m. Then, the mixture was added to cultured cells for infection and incubated for an additional 24 h. The luminescence was measured by Britelite plus Reporter Gene Assay System (PerkinElmer). The 50% inhibitory dilution (IC50) was defined as the antibody concentration at which the relative light units (RLUs) were reduced by 50% compared with the virus control wells (virus + cells) after subtraction of the background RLUs in the control groups with cells only. The IC50 values were calculated with non-linear regression using GraphPad Prism 8 (GraphPad Software, Inc.).

**Cryo-EM data collection and processing**. The prefusion-stabilized Lassa GPC trimer was deposited on a grid unliganded or incubated with 2-fold molar excess per protomer of nanobody D5 and antibody Fab fragments of 8.11G. A volume of 2.3 μl of the complex at 1 mg/ml concentration was deposited on a C-flat grid (protochip.com) and the grid was vitrified using an FEI Vitrobot Mark IV with a wait time of 30 seconds, blot time of 3 seconds and blot force of 1. Data collection was performed on a Titan Krios electron microscope with Leginon (14) with a Gatan K2 Summit direct detection device for the unliganded trimer and a Gatan K3 for the liganded sample. Exposures were collected in movie mode for a 10 s with the total dose of 56.52 e–/Å$^2$ fractionated over 50 raw frames for the unliganded and for a 2 s with the total dose of 51.15 e–/Å$^2$ fractionated over 50 raw frames for the liganded. Images were pre-processed using Appion (15, 16); individual frames were aligned and dose-weighted using MotionCor2 (Zheng et al., 2017). CTFFind4 (17, 18) was used to estimate the CTF and DoG Picker (15, 16) was used for particle picking. RELION (7) was used for particle extraction. CryoSPARC 3.3 (19) was used for 2D classifications, ab initio 3D reconstruction, homogeneous refinement, and nonuniform 3D refinement.

Coordinates from PDB ID 5VK2 was used for initial fit to the reconstructed map. This was followed by simulated annealing and real space refinement in Phenix (21) and then iteratively processed with manual fitting of the coordinates in Coot (22). Geometry and map fitting were evaluated throughout the process using Molprobity (23) and EMRinger (24). PyMOL (www.pymol.org) and ChimeraX were used to generate figures.

**Data and software availability**. Cryo-EM maps and fitted coordinates are in the process of being deposited with database codes EMDB-xxxx, xxxx, xxxx and PDB ID xxxx respectively.

### Table S1

**Table S1. BLI binding data of D5 nanobody and Fabs of human Lassa nAbs toward prefusion-stabilized LASV GPC trimer.**

| Nanobody | | GPC | Fab | | | Fab | | |
|---|---|---|---|---|---|---|---|---|
| B8 | KD (M) | 1.88 E-08 | 10.4B | KD (M) | 6.63 E-08 | 36.1F | KD (M) | 1.52 E-08 |
| | Kon (M$^{-1}$s$^{-1}$) | 1.76 E+05 | | Kon (M$^{-1}$s$^{-1}$) | 5.38 E+04 | | Kon (M$^{-1}$s$^{-1}$) | 1.71 E+05 |
| | Kdis (s$^{-1}$) | 3.31 E-03 | | Kdis (s$^{-1}$) | 3.57 E-03 | | Kdis (s$^{-1}$) | 2.61 E-03 |
| B10 | KD (M) | 4.45 E-08 | 12.1F | KD (M) | 3.64 E-08 | 18.5C | KD (M) | 7.31 E-09 |
| | Kon (M$^{-1}$s$^{-1}$) | 2.23 E+05 | | Kon (M$^{-1}$s$^{-1}$) | 2.81 E+04 | | Kon (M$^{-1}$s$^{-1}$) | 5.35 E+04 |
| | Kdis (s$^{-1}$) | 9.92 E-03 | | Kdis (s$^{-1}$) | 1.02 E-03 | | Kdis (s$^{-1}$) | 3.91 E-04 |
| C3 | KD (M) | 1.97 E-08 | 19.7E | KD (M) | 1.57 E-07 | 25.6A | KD (M) | 1.54 E-08 |
| | Kon (M$^{-1}$s$^{-1}$) | 3.49 E+05 | | Kon (M$^{-1}$s$^{-1}$) | 8.48 E+03 | | Kon (M$^{-1}$s$^{-1}$) | 1.64 E+05 |
| | Kdis (s$^{-1}$) | 6.88 E-03 | | Kdis (s$^{-1}$) | 1.33 E-03 | | Kdis (s$^{-1}$) | 2.53 E-03 |
| D5 | KD (M) | 2.67 E-08 | 8.11G | KD (M) | 3.71 E-08 | 37.7H | KD (M) | 4.47 E-09 |
| | Kon (M$^{-1}$s$^{-1}$) | 1.21 E+05 | | Kon (M$^{-1}$s$^{-1}$) | 3.68 E+04 | | Kon (M$^{-1}$s$^{-1}$) | 1.85 E+05 |
| | Kdis (s$^{-1}$) | 3.23 E-03 | | Kdis (s$^{-1}$) | 1.37 E-03 | | Kdis (s$^{-1}$) | 8.28 E-04 |
| | | | 25.10C | KD (M) | 8.56 E-09 | | | |
| | | | | Kon (M$^{-1}$s$^{-1}$) | 2.82 E+05 | | | |
| | | | | Kdis (s$^{-1}$) | 2.41 E-03 | | | |

### Figure S1

| Construct description | Physiological | Lower pH (5.5) | Higher temp (56° C) |
|---|---|---|---|
| Lassa_50A10THS | 1.4936 | 3.0184 | 1.047 |
| Lassa_50A15THS | 1.4354 | 3.0665 | 0.9952 |
| Lassa_50A12THS | 0.9944 | 2.3964 | 0.6915 |
| Lassa_R325CM359C_deloriDS_ | 0.501 | 0.7142 | 0.3089 |
| Lassa_R207C_E239P_4R_G360C_T331P | 0.3274 | 0.3487 | 0.2654 |
| Lassa_FdTHS | 0.3234 | 0.6245 | 0.1757 |
| Lassa_DS1 | 0.3108 | 0.2583 | 0.1827 |
| Lassa_R207C_E239P_4R_G360C_A343G | 0.2984 | 0.2968 | 0.1912 |
| Lassa_R207C_E239P_4R_G360C_S332G | 0.2873 | 0.3602 | 0.1783 |
| Lassa_R207C_E239P_4R_G360C_S332P | 0.2307 | 0.2678 | 0.1485 |
| **LASSA_5VK2** | 0.2287 | 0.1834 | 0.1508 |
| Lassa_R207C_E239P_4R_G360C_T331G | 0.2285 | 0.2764 | 0.1708 |
| Lassa_303Y | 0.2241 | 0.2088 | 0.1486 |
| Lassa_R207C_E239P_4R_G360C_N342G | 0.2237 | 0.2346 | 0.1776 |
| Lassa_248W | 0.2229 | 0.2029 | 0.1504 |
| Lassa_R207C_E239P_4R_G360C_Q330G | 0.216 | 0.2095 | 0.1491 |
| Lassa_R207C_E239P_4R_G360C_Q330P | 0.2142 | 0.2286 | 0.1609 |
| Lassa_bRFdTHS | 0.2055 | 0.2751 | 0.1181 |
| Lassa_R207C_E239P_4R_G360C_K327G | 0.1892 | 0.1963 | 0.1461 |
| Lassa_R207C_E239P_4R_G360C_K327P | 0.1628 | 0.1784 | 0.1214 |
| Lassa_305W | 0.1587 | 0.1298 | 0.102 |
| Lassa_303W | 0.1542 | 0.1713 | 0.143 |
| LassaGP_144C147C | 0.1524 | 0.1539 | 0.1115 |
| LassaGP_81C319C | 0.1518 | 0.1253 | 0.1337 |
| Lassa_R207C_E239P_4R_G360C_V341G | 0.1517 | 0.1312 | 0.0987 |
| Lassa_305F | 0.1344 | 0.1516 | 0.1118 |
| LassaGP_87C198C | 0.1274 | 0.1107 | 0.1078 |
| Lassa_R193M_Q247M_K339M | 0.1271 | 0.1199 | 0.107 |
| Lassa_R193M_D211M_Q247M | 0.1221 | 0.0983 | 0.0969 |
| Lassa_Q247M_K339M | 0.116 | 0.1341 | 0.0982 |
| Lassa_R193M_D211M_K339M | 0.1102 | 0.1034 | 0.0967 |
| Lassa_R207C_E239P_4R_G360C_N346G | 0.106 | 0.1248 | 0.0953 |
| LassaGP_113C165C | 0.1056 | 0.1165 | 0.1005 |
| Lassa_D211M_Q247M_K339M | 0.1033 | 0.0996 | 0.1093 |
| Lassa_R193M_D211M_Q247M_K339M | 0.1012 | 0.0909 | 0.0894 |
| LassaGP_246C347C | 0.0993 | 0.087 | 0.0954 |
| LassaGP_356C361C | 0.0966 | 0.0917 | 0.086 |
| LassaGP_196C240C | 0.0961 | 0.105 | 0.123 |
| LassaGP_168C184C | 0.096 | 0.0739 | 0.0732 |
| LassaGP_348C343C | 0.096 | 0.0904 | 0.1082 |
| Lassa_M73CI286C | 0.0933 | 0.0926 | 0.1343 |
| Lassa_DiSu1_R193M_D211M_Q247M_R250F_K339M | 0.0915 | 0.0958 | 0.1004 |
| LassaGP_1 | 0.0914 | 0.0859 | 0.0884 |
| LassaGP_197C234C | 0.0912 | 0.0803 | 0.0919 |
| LassaGP_2 | 0.0905 | 0.0788 | 0.0907 |
| Lassa_L326CG208GC | 0.0901 | 0.0894 | 0.0912 |
| LassaGP_348C260C | 0.0896 | 0.0732 | 0.0815 |
| Lassa_403W | 0.0895 | 0.1111 | 0.0878 |
| **LASSA WT** | 0.0894 | 0.1118 | 0.0896 |
| Lassa_R193M_D211M_Q247M_R250F_K339M | 0.0886 | 0.0864 | 0.0914 |
| Lassa_R207C_E239P_4R_G360C_L344G | 0.0882 | 0.0759 | 0.0816 |
| LassaGP_3 | 0.0871 | 0.0829 | 0.1431 |
| LassaGP_107C217C | 0.085 | 0.0938 | 0.1188 |
| Lassa_I345P | 0.0841 | 0.0764 | 0.0809 |
| Lassa_R207C_E239P_4R_G360C_I345G | 0.0817 | 0.0904 | 0.0943 |
| Lassa_N74CL285C | 0.0812 | 0.0824 | 0.0846 |
| LassaGP_85C240C | 0.0812 | 0.0722 | 0.0794 |
| Lassa_T263CA343C | 0.0809 | 0.073 | 0.0867 |
| LassaGP_143C244C | 0.0807 | 0.0742 | 0.0792 |
| LassaGP_198C234C | 0.0805 | 0.0771 | 0.0823 |
| Lassa_396W | 0.0803 | 0.0819 | 0.0844 |
| LassaGP_62C407C | 0.0787 | 0.0642 | 0.0744 |
| Lassa_407W | 0.0786 | 0.0861 | 0.0778 |
| LassaGP_353C363C | 0.078 | 0.0804 | 0.0772 |
| **LASSA GP2** | 0.0769 | 0.0685 | 0.0811 |
| Lassa_Q69CY371CR325CM359C | 0.0764 | 0.0645 | 0.0758 |
| Lassa_N74CM284C | 0.0759 | 0.07 | 0.0773 |
| LassaGP_314C344C | 0.0759 | 0.0675 | 0.0814 |
| LassaGP_311C345C | 0.0748 | 0.07 | 0.077 |
| Lassa_DiSu2_R193M_D211M_Q247M_R250F_K339M | 0.0741 | 0.0818 | 0.0847 |
| Lassa_DiSu3_R193M_D211M_Q247M_R250F_K339M | 0.0738 | 0.0689 | 0.0808 |
| **no DNA** | 0.0736 | 0.0662 | 0.0772 |
| Lassa_DS4 | 0.0729 | 0.0673 | 0.0881 |
| Lassa_DS2 | 0.0727 | 0.0682 | 0.0799 |
| LassaGP_369C383C | 0.0723 | 0.0723 | 0.0759 |
| LassaGP_362C389C | 0.072 | 0.0704 | 0.075 |
| Lassa_DiSu1_2_R193M_D211M_Q247M_R250F_K339M | 0.0719 | 0.0665 | 0.0808 |
| LassaGP_145C251C | 0.0719 | 0.0656 | 0.0733 |
| LassaGP_347C260C | 0.0718 | 0.064 | 0.1002 |
| **no DNA** | 0.0717 | 0.0647 | 0.0727 |
| Lassa_DiSu1_3_R193M_D211M_Q247M_R250F_K339M | 0.0712 | 0.0664 | 0.0795 |
| Lassa_DiSu1_2_3_R193M_D211M_Q247M_R250F_K339M | 0.0712 | 0.0667 | 0.0778 |
| LassaGP_356C363C | 0.0707 | 0.0635 | 0.0731 |
| LassaGP_368C386C | 0.0705 | 0.0649 | 0.0859 |
| Lassa_DS3 | 0.0705 | 0.0691 | 0.0792 |
| Lassa_E392CG98C | 0.0701 | 0.0649 | 0.0774 |
| LassaGP_166C220C | 0.0699 | 0.0634 | 0.0778 |
| **LASSA GP1** | 0.0683 | 0.0626 | 0.0734 |

| Construct description | Physiological | Lower pH (5.5) | Higher temp (56° C) |
|---|---|---|---|
| Lassa_FD2 | 0.4772 | 1.002 | 0.4313 |
| Lassa_D347P | 0.4744 | 0.6894 | 0.396 |
| Lassa_FD3 | 0.4055 | 0.8205 | 0.3533 |
| Lassa_FD2F305 | 0.3608 | 0.74 | 0.263 |
| Lassa_FD4 | 0.3357 | 0.5328 | 0.2846 |
| Lassa_FD1 | 0.2582 | 0.5977 | 0.2527 |
| **LASSA_5VK2** | 0.2119 | 0.2572 | 0.1772 |
| Lassa_MF1 | 0.2079 | 0.27 | 0.1932 |
| Lassa_MF3 | 0.1626 | 0.2868 | 0.1779 |
| Lassa_FD2W303 | 0.1515 | 0.2317 | 0.1536 |
| Lassa_R193M | 0.147 | 0.1659 | 0.1182 |
| Lassa_R193K339E | 0.1462 | 0.1643 | 0.134 |
| Lassa_K339M | 0.1424 | 0.1397 | 0.1163 |
| Lassa_MF2 | 0.1334 | 0.2285 | 0.1421 |
| Lassa_Q247M | 0.1314 | 0.1668 | 0.1181 |
| Lassa_1nza | 0.1296 | 0.2481 | 0.1659 |
| Lassa_R250F | 0.1289 | 0.1633 | 0.1239 |
| Lassa_R193K339D | 0.122 | 0.1562 | 0.1151 |
| Lassa_R193M_K339M | 0.1218 | 0.1346 | 0.1114 |
| LassaSS4 | 0.1205 | 0.1081 | 0.1051 |
| Lassa_R193M_Q247M | 0.1169 | 0.1336 | 0.1109 |
| LassaSS5 | 0.1159 | 0.0796 | 0.0846 |
| Lassa_MF4 | 0.1156 | 0.1656 | 0.1306 |
| Lassa_D211M_Q247M | 0.1093 | 0.1321 | 0.1119 |
| Lassa_FD2C72C368 | 0.1072 | 0.0887 | 0.0885 |
| Lassa_D211M | 0.1066 | 0.1208 | 0.1018 |
| Lassa_Q189LR193L_M351F | 0.105 | 0.1137 | 0.0984 |
| Lassa_b_v6 | 0.1027 | 0.1101 | 0.0848 |
| Lassa_MF4C67C373 | 0.1026 | 0.0862 | 0.0805 |
| Lassa_N346P | 0.0989 | 0.0682 | 0.0799 |
| Lassa_342F_N346FV388L | 0.0975 | 0.1026 | 0.0952 |
| Lassa_R193M_D211M | 0.0958 | 0.1003 | 0.0964 |
| Lassa_D211M_K339M | 0.0926 | 0.0968 | 0.0886 |
| LassaSS8 | 0.0898 | 0.0909 | 0.0959 |
| Lassa_Disu1_3 | 0.0881 | 0.0689 | 0.0777 |
| Lassa_DiSu3 | 0.086 | 0.0913 | 0.1078 |
| Lassa_C263C352P130 | 0.0849 | 0.0753 | 0.103 |
| **LASSA WT** | 0.0823 | 0.0809 | 0.0771 |
| Lassa_MF2sc1 | 0.0799 | 0.1071 | 0.0875 |
| Lassa_Disu1_2_3 | 0.0794 | 0.0771 | 0.0802 |
| Lassa_FD3sc1 | 0.0779 | 0.1014 | 0.0894 |
| LassaSS6 | 0.0767 | 0.0861 | 0.0901 |
| Lassa_D306FD302F | 0.0766 | 0.0753 | 0.0782 |
| Lassa_C138C254 | 0.0765 | 0.0768 | 0.0946 |
| **LASSA GP1** | 0.0762 | 0.0659 | 0.0768 |
| Lassa_MF4C72C368 | 0.0757 | 0.0844 | 0.0863 |
| Lassa_T70CW370C | 0.0753 | 0.079 | 0.0881 |
| Lassa_MF3sc1 | 0.0747 | 0.0771 | 0.0807 |
| Lassa_C263C352_C138C254 | 0.0746 | 0.0701 | 0.0836 |
| Lassa_b_v5 | 0.0746 | 0.0801 | 0.089 |
| Lassa_L71CY369C | 0.0738 | 0.1221 | 0.0889 |
| Lassa_FD2sc1 | 0.0736 | 0.085 | 0.0887 |
| Lassa_C263C352sc1 | 0.0732 | 0.0798 | 0.0866 |
| Lassa_turn329v2 | 0.0725 | 0.0768 | 0.0806 |
| Lassa_FD2sc3 | 0.0722 | 0.0716 | 0.0853 |
| Lassa_C263C348 | 0.0722 | 0.0735 | 0.0841 |
| Lassa_C263C352sc2 | 0.072 | 0.0766 | 0.0787 |
| Lassa_turn329v3 | 0.0717 | 0.0809 | 0.0811 |
| Lassa_turn170_v1 | 0.0712 | 0.0677 | 0.0768 |
| Lassa_342F_N346F | 0.0712 | 0.0658 | 0.0788 |
| Lassa_FD2C343C348 | 0.0708 | 0.0883 | 0.0834 |
| Lassa_C263C352P131 | 0.0707 | 0.069 | 0.0774 |
| Lassa_D306LD302L | 0.0706 | 0.0735 | 0.0749 |
| **LASSA GP2** | 0.0706 | 0.0708 | 0.0942 |
| Lassa_FD2C67C373 | 0.0704 | 0.0675 | 0.0785 |
| Lassa_turn170_v2 | 0.0702 | 0.0687 | 0.0801 |
| Lassa_b_v7 | 0.0702 | 0.066 | 0.0761 |
| Lassa_b_v2 | 0.07 | 0.0679 | 0.0758 |
| Lassa_C263C349 | 0.0698 | 0.0691 | 0.0859 |
| Lassa_Disu2_3 | 0.0698 | 0.0694 | 0.0776 |
| Lassa_FD2C263C352sc1 | 0.0696 | 0.0698 | 0.0823 |
| Lassa_MF4C263C352sc1 | 0.0696 | 0.0671 | 0.0782 |
| Lassa_385s | 0.0694 | 0.0716 | 0.0776 |
| Lassa_FD2sc2 | 0.0693 | 0.0702 | 0.0845 |
| Lassa_DSS1E72CK368CQ69CY371C | 0.0692 | 0.0653 | 0.0761 |
| **no DNA** | 0.0689 | 0.0654 | 0.073 |
| Lassa_b_v4 | 0.0688 | 0.0679 | 0.074 |
| Lassa_b_v1 | 0.0686 | 0.0652 | 0.0756 |
| Lassa_E72CK368C | 0.0684 | 0.0695 | 0.085 |
| Lassa_MF4C263C352 | 0.0683 | 0.0673 | 0.0838 |
| Lassa_DiSu1_2 | 0.0682 | 0.0799 | 0.0782 |
| Lassa_b_v3 | 0.0681 | 0.0667 | 0.0768 |
| Lassa_I410E_M414E | 0.068 | 0.0644 | 0.0752 |
| Lassa_Q69CY371C | 0.0678 | 0.0689 | 0.0769 |
| **no DNA** | 0.0678 | 0.0679 | 0.074 |
| Lassa_turn329v4 | 0.0676 | 0.0669 | 0.0767 |
| Lassa_FD2C263C352 | 0.067 | 0.0675 | 0.0783 |
| Lassa_C263C352C67C373 | 0.0658 | 0.0657 | 0.0735 |

**Fig. S1. Antigenic screening of designed prefusion-stabilized Lassa GPC constructs.**
We designed 164 variants, which we screened for binding to the quaternary-specific GPC antibody 37.7H under physiological conditions, at lower pH (5.5) and at higher temperature (56° C). Numbers indicate ELISA results ($OD_{450}$) for recognition by 37.7H. Top hits for each of the two plates are highlighted in red fonts. Controls are indicated with green background. Highlighted in blue in the “Construct description” column are the two designs, inter-protomer disulfide and foldon fusion, that were combined in the final construct for detailed characterization in this study after evaluation of their behavior at 1 liter expression and the ability to combine them.

### Figure S2

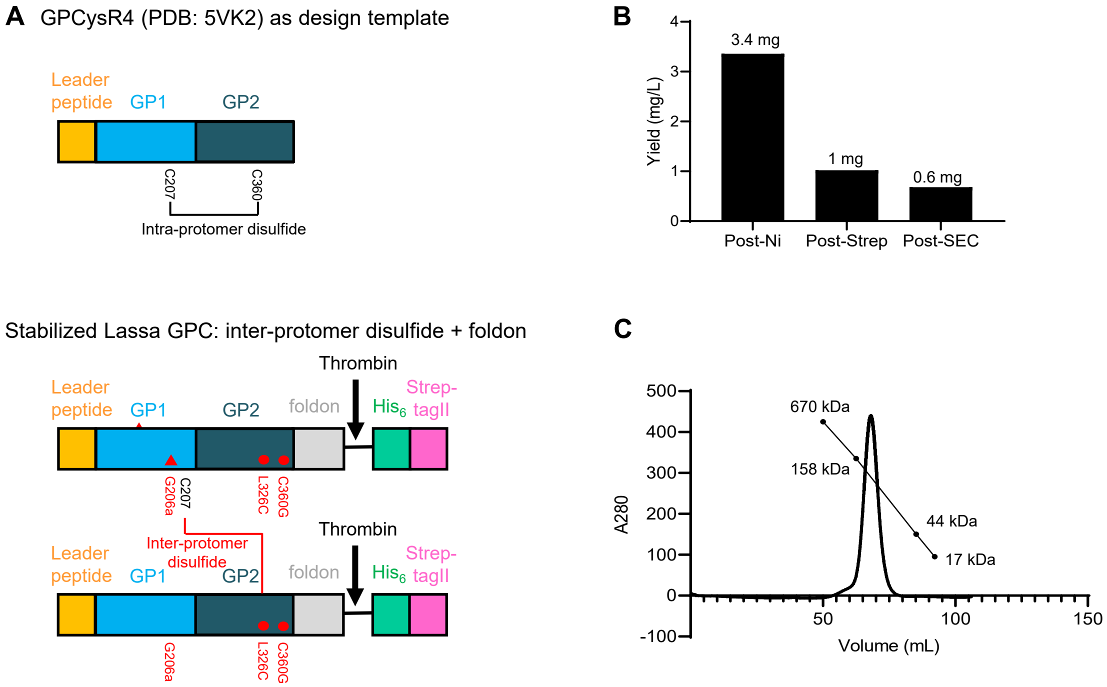


**Fig. S2. Construct, yield, and purification profile of stabilized soluble Lassa GPC trimer.**

(A) Schematic showing the design of GPCysR4 (top) and stabilized soluble Lassa GPC trimer (below). The original C207-C360 intra-protomer disulfide in GPCysR4 was abolished by mutating C360G. New inter-protomer disulfide was created between C207 and L326C. Insertion of G206a allowed optimal geometry for such disulfide bond formation. T4-fibritin (foldon) trimerization domain was also introduced at the C-terminus to fix the base of the trimer.

(B) Protein yield of the stabilized soluble GPC trimer following nickel-affinity (Ni), streptavidin-affinity (Strep), and size exclusion (SEC) purification.

(C) SEC profile of the stabilized soluble GPC trimer on Superdex 200 16/600 column.

### Figure S3

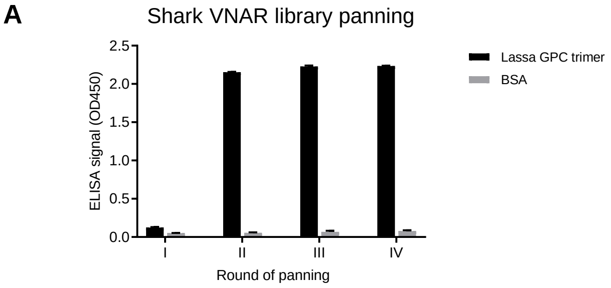


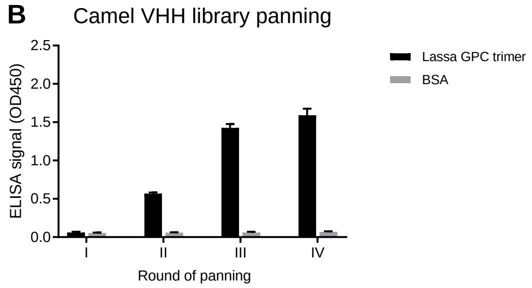


| Round | Input phage (Colony forming unit, CFU) | Output phage (Colony forming unit, CFU) |
|---|---|---|
| I | $1\times10^{12}$ | $5\times10^{3}$ |
| II | $1\times10^{12}$ | $5\times10^{3}$ |
| III | $1\times10^{12}$ | $5\times10^{5}$ |
| IV | $1\times10^{12}$ | $2\times10^{6}$ |

| Round | Input phage (Colony forming unit, CFU) | Output phage (Colony forming unit, CFU) |
|---|---|---|
| I | $1\times10^{12}$ | $5\times10^{3}$ |
| II | $1\times10^{12}$ | $3\times10^{2}$ |
| III | $1\times10^{12}$ | $1\times10^{5}$ |
| IV | $1\times10^{12}$ | $6\times10^{6}$ |

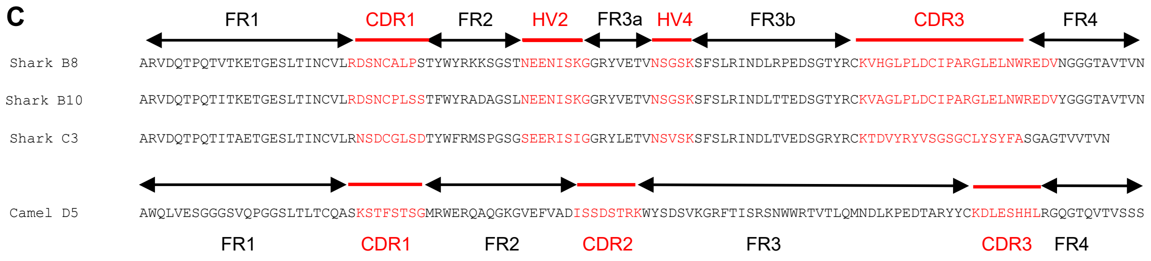


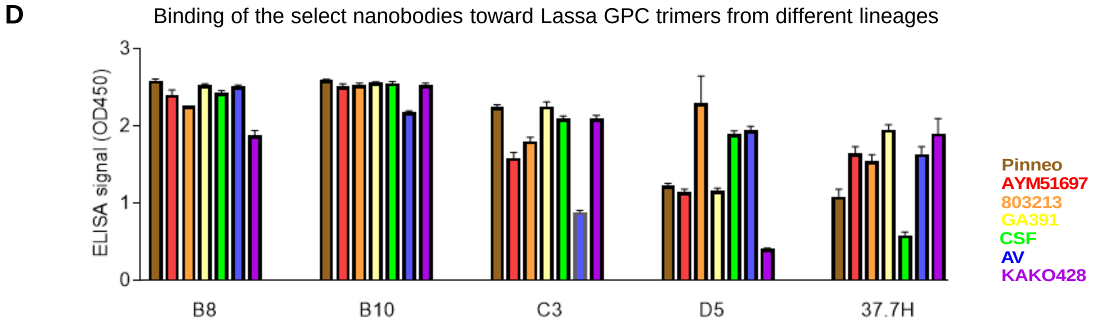


**Fig. S3. Identification of nanobodies by panning shark VNAR and camel VHH libraries against stabilized soluble Lassa GPC trimer.**

(A) Phage ELISA following different rounds of stabilized Lassa GPC panning by the shark VNAR library. Input phage and output phage titers were tabulated below.

(B) Phage ELISA following different rounds of stabilized Lassa GPC panning by the camel VHH library. Input phage and output phage titers were tabulated below.

(C) Sequences and IMGT annotation of the nanobodies.

(D) Binding of the select nanobodies toward trimers from 7 different lineages of Lassa virus.

Figure S4

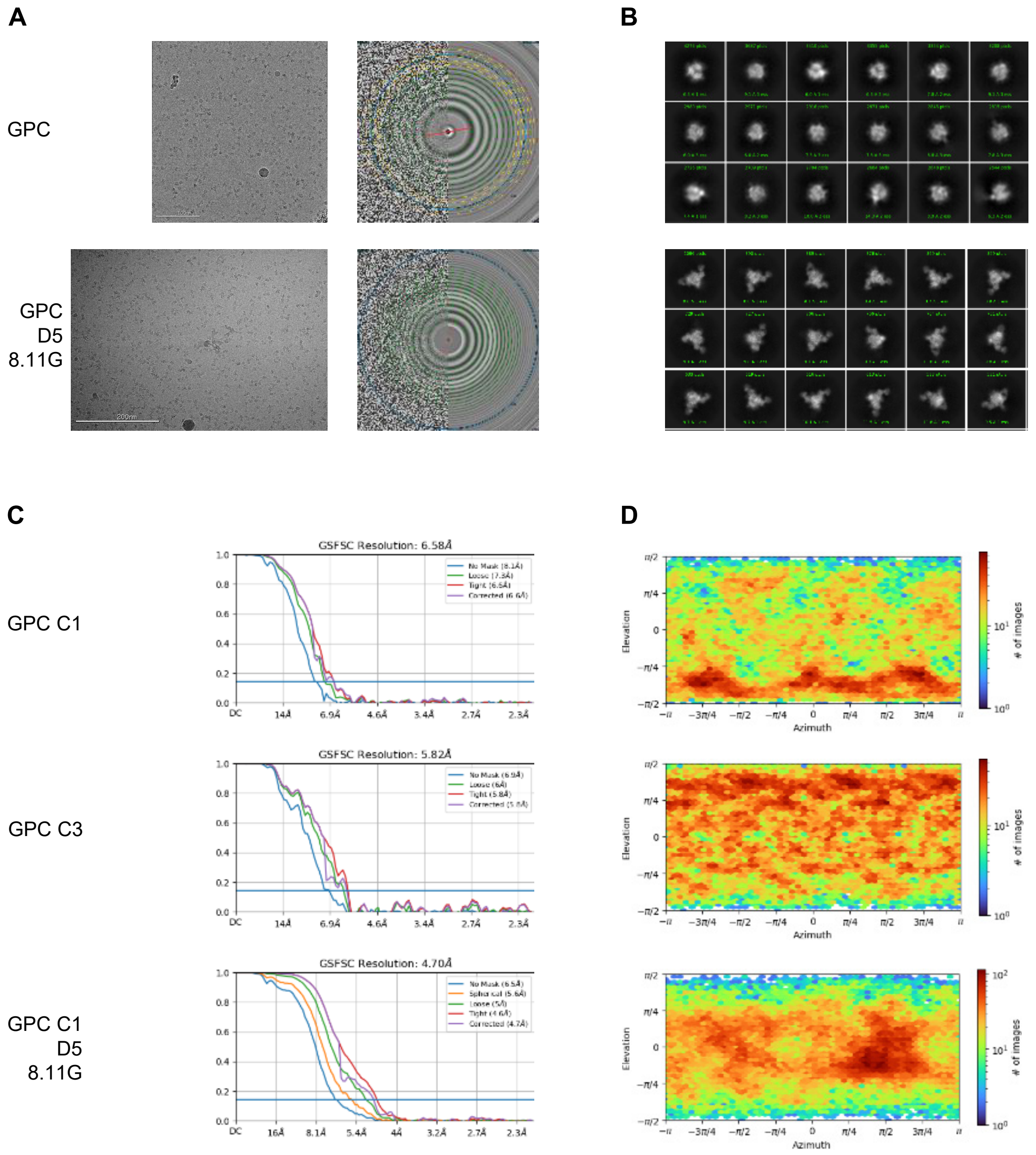


**Fig. S4. Cryo-EM details of prefusion-stabilized GPC trimer and in complex with human Fab 8.11G and nanobody D5.**
(A) Representative micrographs and CTFs of the micrographs are shown.
(B) Representative 2D class averages are shown.
(C) The gold-standard Fourier shell correlation are shown with the resolution for the three maps.
(D) The orientations of all particles used in the final refinements are shown as heatmaps.
